# Non-Linear Realignment Improves Hippocampus Subfield Segmentation

**DOI:** 10.1101/597856

**Authors:** Thomas B Shaw, Steffen Bollmann, Nicole T Atcheson, Christine Guo, Jurgen Fripp, Olivier Salvado, Markus Barth

## Abstract

Participant movement can deleteriously affect MR image quality. Further, for the visualization and segmentation of small anatomical structures, there is a need to improve image quality, specifically signal-to-noise ratio (SNR) and contrast-to-noise ratio (CNR), by acquiring multiple anatomical scans consecutively. We aimed to ameliorate movement artefacts and increase SNR in a high-resolution turbo spin-echo (TSE) sequence acquired thrice using non-linear realignment in order to improve segmentation consistency of the hippocampus subfields. We assessed the method in young healthy participants, Motor Neurone Disease patients, and age matched controls. Results show improved image segmentation of the hippocampus subfields when comparing template-based segmentations with individual segmentations with Dice overlaps N=51; *ps* < 0.001 (Friedman’s test) and higher sharpness *ps* < 0.001 in non-linearly realigned scans as compared to linearly, and arithmetically averaged scans.

## 1. Introduction

Imaging the subfields of the hippocampal formation with Magnetic Resonance Imaging (MRI) has garnered great interest in recent years. Characterising subfield tissue may elucidate susceptibility to – and provide more sensitive biomarkers for - neurodegenerative diseases including Alzheimer’s disease (AD) (Balachandar et al., 2015; Boutet et al., 2014; Henry et al., 2011; Jacobsen et al., 2017; Kerchner et al., 2012; La Joie et al., 2013; Maruszak & Thuret, 2014; Pluta, Yushkevich, Das, & Wolk, 2012), as the subfields contain different cell types and functional properties (Duvernoy, Cattin, Risold, Vannson, & Gaudron, 2013). Increasing spatial resolution for MRI through improved acquisition and/or post-processing techniques improves segmentation of various tissue types or anatomical structures (Thomas et al., 2008). Consequently, there is a need to improve signal-to-noise ratio (SNR) of scans by going to higher field strengths (ultra-high field [UHF] MRI, 7T and above) and by acquiring multiple repetitions of the same scan in a single session, which leads to longer acquisition times.

Previous UHF *in vivo* hippocampus subfield segmentation studies (for review, see Giuliano et al., 2017) have used dedicated T2-weighted Turbo-Spin Echo (TSE), Gradient Echo, or multi-echo sequences that, due to multiple refocusing pulses, exhibit differing contrast signals and intensity characteristics for different tissue classes, and consequently the laminae of the hippocampus (Marques & Norris, 2018; Winterburn et al., 2013). With higher field strengths and better coils comes the possibility of increasing the number of repetitions due to shorter scan times per acquisition. We have previously utilised three repetitions of a four minute TSE sequence (12 minutes total acquisition time) that cover the whole hippocampus with a high in-plane resolution of 0.4 mm^2^ x 0.4 mm^2^ and a 0.8 mm^2^ slice thickness. These acquisitions were combined to yield a minimum deformation average model for investigating hippocampus subfields (Jacobsen et al., 2017). The sequence was designed to be repeated thrice to both a) boost SNR and b) to be able to potentially discard one of the repetitions in case of participant movement. Unfortunately, finding the optimal realignment between these anatomical scans can be difficult due to gradient nonlinearities (Reuter, Rosas, & Fischl, 2010) especially at ultra-high field. Reuter et al. (2010) used rigid registrations for realignment of anatomical T1w scans in an intermediate space as rigid parts of the brain and skull are assumed to go unchanged in the short time between scan repetitions (save for rigid location changes). At higher fields strengths and resolutions, noise from movement interacts with gradient nonlinearities and may cause non-linear distortions in images unable to be corrected by affine registrations alone.

Participant movement is more likely for long MRI scans and can deteriorate image quality (Kochunov et al., 2006), potentially leading to unusable scans due to movement artefacts, and subsequent increased costs. Further, patients with neurodegenerative disorders, elderly, young, and highly anxious participants, have a high propensity for movement in the scanner (Yushkevich et al., 2015). Previous work examining the benefits of motion correction in anatomical MRI have used either prospective or retrospective realignment techniques. Prospective techniques (Maclaren, Herbst, Speck, & Zaitsev, 2013) mainly utilise navigators and motion tracking devices to correct for motion online. Retrospective realignment techniques may take magnitude data and attempt to estimate transformations between images to attain good registrations between them (e.g., Reuter et al., 2010), similar to fMRI realignment.

In human neuroimaging with a supine patient position, rotation commonly occurs at the posterior of the head, resulting in rotation-related blurring and movement artefacts progressing in severity anteriorly (Maclaren et al., 2013). Blurring and ghosting artefacts are somewhat prevalent in imaging the hippocampus *in vivo,* which has tightly packed laminae and multiple tissue contrast signals that blur together causing partial volume effects. This motion is inherent to the longer acquisition times necessary to capture the structure of the hippocampus (Marrakchi-Kacem et al., 2016). Mitigating movement artefacts in image post-processing is therefore an important challenge to overcome in neuroimaging.

Previous work (Bollmann, Bollmann, Puckett, Janke, & Barth, 2017) has shown that fMRI realignment can be completed using an iterative averaging model (or template) to account for movement in participants and decrease partial volume effects. We aim to utilise a similar approach on anatomical scans with high resolution in order to reduce partial volume effects and increase SNR. Due to the small size of the hippocampus and its laminar structure, partial volume effects can lead to misclassification of subfields. To ameliorate movement artefacts, boost SNR and sharpness, and improve image segmentation reliability and validity, we implemented a retrospective realignment technique that iteratively estimates non-linear transformations between multiple images to fit to an evolving model of anatomical consistency to attain improved segmentation results.

In the present study, quantitative metrics for examination of the most effective registration technique were chosen based in part on the work of Fonov and Collins (2018). It was determined that, much like in the original MP2RAGE paper (Marques et al., 2010), a useful metric for examining the effectiveness of a registration technique was through measuring segmentation performance and consistency (as qualitative analysis of motion correction is not sufficient). Labelling the subfields of the hippocampus with automatic segmentation strategies is now feasible with software including Automatic Segmentation of Hippocampal Subfields (ASHS; Yushkevich et al., 2015), and Freesurfer (Iglesias et al., 2015) all able to segment hippocampal subfields of conventional T1w and dedicated T2w TSE scans in multi-contrast approaches.

We aimed to explore the effectiveness of our technique in healthy controls (HCs) and patients with Motor Neurone Disease (MND) by assessing registration consistency. Intuitively, image registration algorithms are more successful in images with more refined details, higher sharpness, and better SNR. Therefore, greater registration consistency will occur between participants if the registration procedure is more effective. If the technique is robust to movement, between-subject registrations will converge more readily due to increased boundary delineation and SNR increases. Thus, we hypothesised that each participant’s non-linearly realigned (1: *Non-Linear*) registration procedure would produce higher segmentation consistency than a linear procedure (similar to fMRI realignment; 2: *Linear*) or a simple averaging procedure (3: *Average*). This is based on the assumption that automatic segmentation algorithms rely on high SNR/CNR and good boundary delineation to function reliably. Finally, due to reduced movement artefacts and higher SNR after processing, we expect that non-linear realignment would produce the sharpest images compared to linear realignment or simple averaging.

## 2. Methods

Using a 7 T whole-body research scanner (Siemens Healthcare, Erlangen, Germany), with maximum gradient strength of 70 mT/m and a slew rate of 200 mT/m/s and a 7T Tx/32 channel Rx head array (Nova Medical, Wilmington, MA, USA) we acquired a 2D TSE sequence (Siemens WIP tse_UHF_WIP729C. Variant: tse2d1_9, TR: 10300ms, TE: 102ms, FA: 132°, FoV: 220mm, voxel size of 0.4 x 0.4 x 0.8mm^3^ Turbo factor of 9; iPAT (GRAPPA) factor 2) thrice of a slab aligned orthogonally to the hippocampus in three ‘groups’: 11 patients diagnosed with MND (Age, M=59.36, SD=7.65), 11 age-matched control participants (HCs, M=60.23, SD=7.65), and 29 young healthy participants (YHPs, M=26.31, SD=0.66) for a total of 51 participants in order to test the robustness of the realignment on a wide range of participants in terms of age (reflected by a different brain anatomy) and movement probability (patient vs. healthy). An anatomical whole-brain T1w was acquired using a prototype MP2RAGE sequence (WIP 900) (Marques et al., 2010; O’Brien et al., 2014) at 0.9mm isotropic voxel size was also acquired (TR/TE/TIs = 4300ms / 2.5ms / 840ms, 2370 ms).

All TSEs were pre-processed by resampling to 0.3mm isotropic, bias field corrected using N4 (Tustison et al., 2010), skull stripped using the co-registered T1w as an initial mask after usage of ROBEX (Iglesias, Liu, Thompson, & Tu, 2011), and intensity normalised between two percent-critical thresholds using NiftiNorm (https://github.com/thomshaw92/nifti_normalise), a Nifti implementation of mincnorm from the medical imaging network common data toolkit (Vincent et al., 2016). To assess registration and segmentation consistency, we tested three different approaches (or ‘methods’; Figure 1): we performed no additional realignment and averaged the slabs arithmetically using ANTs AverageImage (1. *Average)*. We also registered TSEs linearly by: i) concatenating images in time using FSL (Jenkinson, Beckmann, Behrens, Woolrich, & Smith, 2012), ii) estimating affine registrations between each image using FSL’s MCFLIRT (Jenkinson, Bannister, Brady, & Smith, 2002) and iii) averaging the image in the time dimension (2. *Linear*). Third, we performed non-linear registrations using ANTs’ (Avants, Tustison, & Song, 2010) SyN (Avants, Epstein, Grossman, & Gee, 2008) registration (Figure 1), with the three TSE scans being iteratively deformed using antsMultivariateTemplateConstruction2.sh into a minimum deformation average (MDA) template containing only anatomically consistent features (3. *Non-Linear*). This method involves averaging the TSE scans to an intermediate space, and registering the individual TSEs to the intermediate space. The average of these transformations is applied to the intermediate template and the model is updated iteratively. We used the default settings and three iterations for the non-linear registrations. We deactivated the ANTs laplacian sharpness filter for fair comparison between realignment methods.

**Figure 1.**
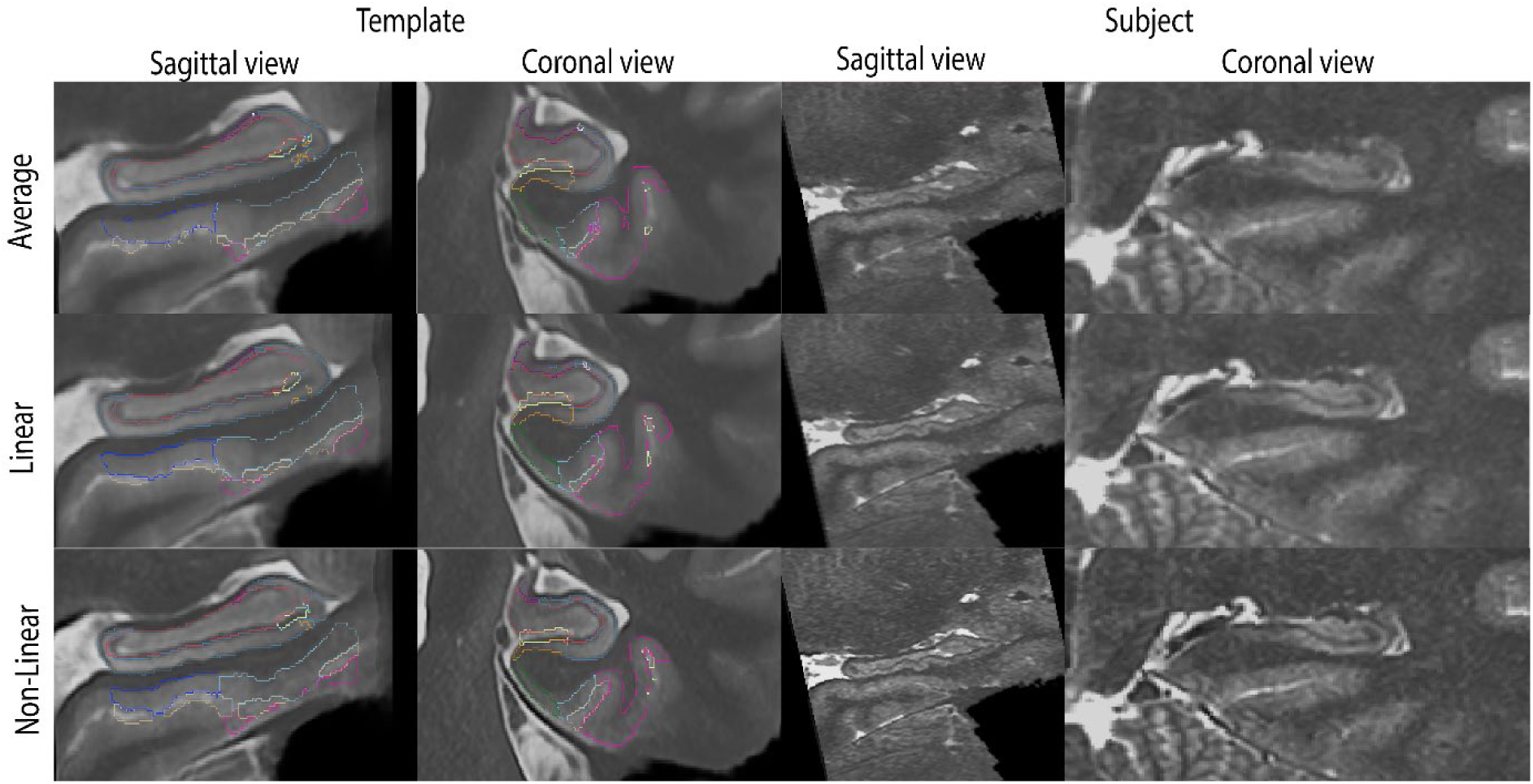
Sagittal and coronal views of the alignment in the hippocampus after the three realignment procedures (Non-Linear, Linear, Average) for the method-templates (left, labelled) and a representative YHP subject that benefited from the non-linear realignment technique (right). Views capture the left hippocampus, and both hippocampi for coronal single-subject view. Edges of the template ASHS segmentations have been superimposed as coloured lines.

Individual YHPs, HCs and MNDs for each method were independently co-registered to a group-and-method-specific MDA template using ANTs to yield nine group-and-method templates (e.g., MND-*Linear*, HC-*Non-Linear*, YHP-*Average,* etc.). Each template was constructed with identical parameters. These nine group-and-method templates and each individual’s realigned TSE (for each method) and their corresponding MP2RAGE scan were labelled using ASHS, which requires both T1w and T2w inputs, (Yushkevich et al., 2015) in native space then warped to their common group-method template space (Figure 1). Segmentation consistency was derived by examining Dice overlaps of segmentation labels between group-method template labels and individual volume labels (subject-to-template overlaps). Sharpness of images was measured to describe brain structure delineation by calculating the median of the derivative of a Gaussian applied to the TSE at 1mm FWHM as described in Fonov and Collins (2018). We provide all the code for this project at https://github.com/thomshaw92/NonLinRegImprovesSegAcc.

For validation, all TSEs were rated manually using adapted methods from Backhausen et al. (2016) and Jones and Marietta (2012), with the criteria of i) Image sharpness (including artefacts), ii) Ringing, iii) Contrast-to-Noise Ratio (CNR; subcortical structures), and 4) CNR (GM and WM). Two raters (NTA: senior research radiographer with 30 years medical imaging experience, and TBS: six years research experience in medical imaging) rated all participants on a scale from 1-3, with 1=pass, 2=check, and 3=fail for quality assurance of scans and to assess motion artefacts before post-processing. A weighted average that favoured ratings of sharpness and subcortical CNR was used to summarise the findings. Intraclass correlation (ICC) estimates and their 95% confidence intervals were calculated using SPSS Statistics Package Version 25 (SPSS Inc, Chicago, IL) based on a mean-rating (*k*=2), absolute agreement, 2-way mixed-effects model.

## 3. Results

Figure 2 shows a single-subject example of registration results for the TSEs. Sharper edges and more subfield information is available in the *Non-Linear* method, followed by *Linear*, and no information available in *Average*. When there is limited movement, small differences can be observed between the three methods.

**Figure 2.**
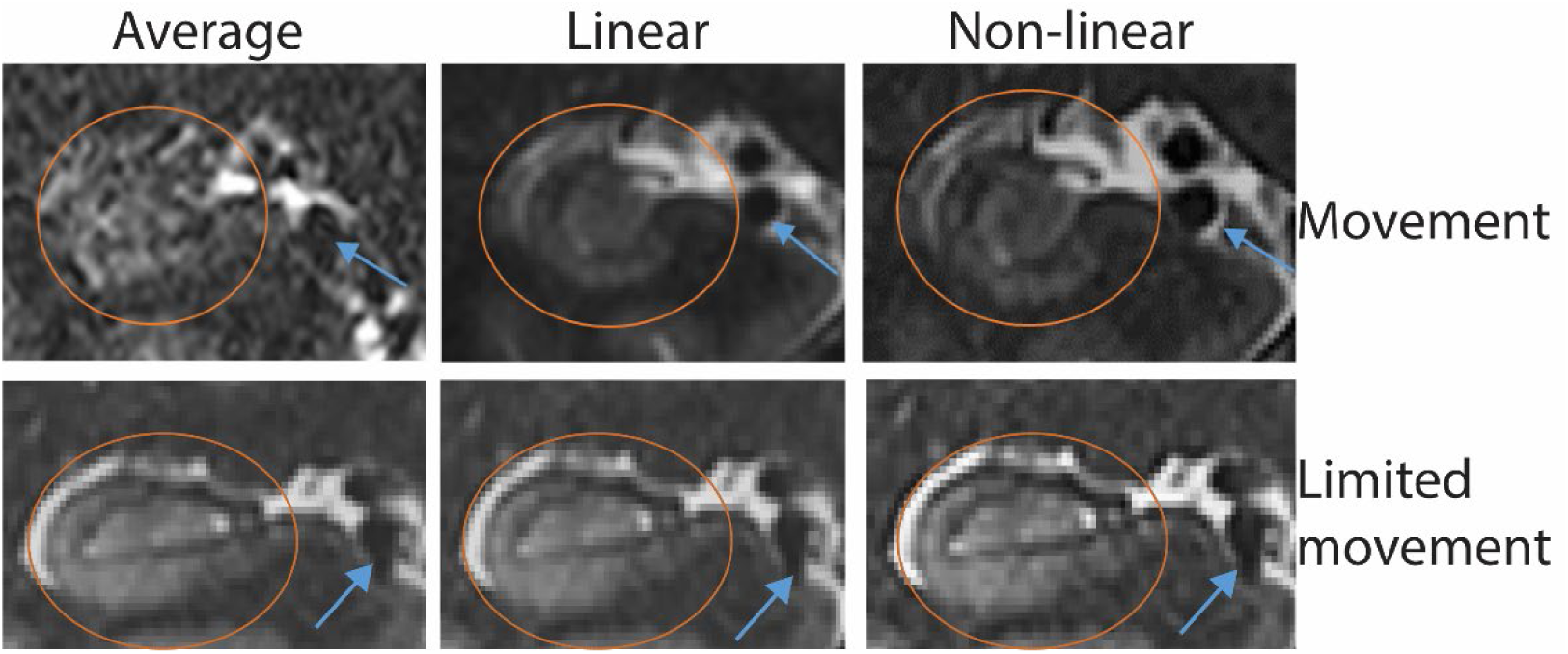
Left to right: Average, Linear, and Non-Linear realignment examples from single participants with high movement (top) and limited movement (bottom) with ellipses over the right hippocampus (coronal plane); blue arrows denote a common vessel.

Collapsing across groups, Dice overlap scores (Figure 3) were found to be significantly different between all groups, (Friedman test *p* < 0.001). Significantly higher overlaps for *Non-Linear* with its method template were found compared to both *Linear* and *Average,* and comparing *Linear* to *Average*, independently (N=51; ** = *ps* < 0.001, Wilcoxon rank sum tests).

**Figure 3.**
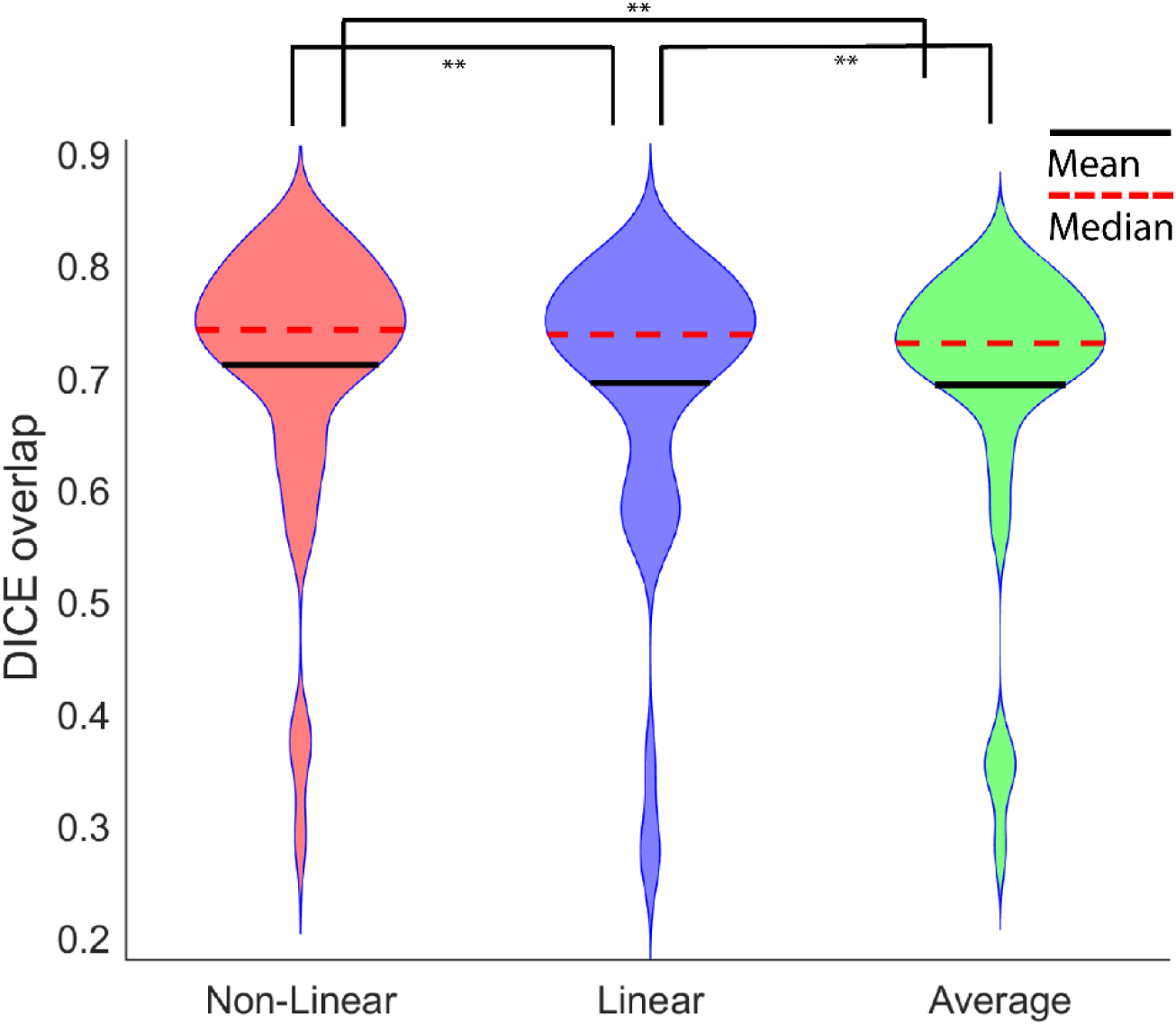
Violin plot for the three realignment methods (Non-Linear, Linear, Average) of Dice overlap scores between individual subject TSEs and its respective method-template, collapsed over the three groups (MND, HC, YHP). ** = ps < 0.001, Wilcoxon rank sum tests).

Sharpness (Figure 4) was significantly higher for *Non-Linear* compared to *Linear* and *Averaging* (*p* < 0.001, Friedman test). Significant differences were found between the three registration techniques, with *Non-Linear* performing better than *Linear,* and A*veraging*, independently (* = *p*s < 0.05). There was no significant difference between *Average* and *Linear* realignment (*p* = .982, Wilcoxon rank sum test).

**Figure 4.**
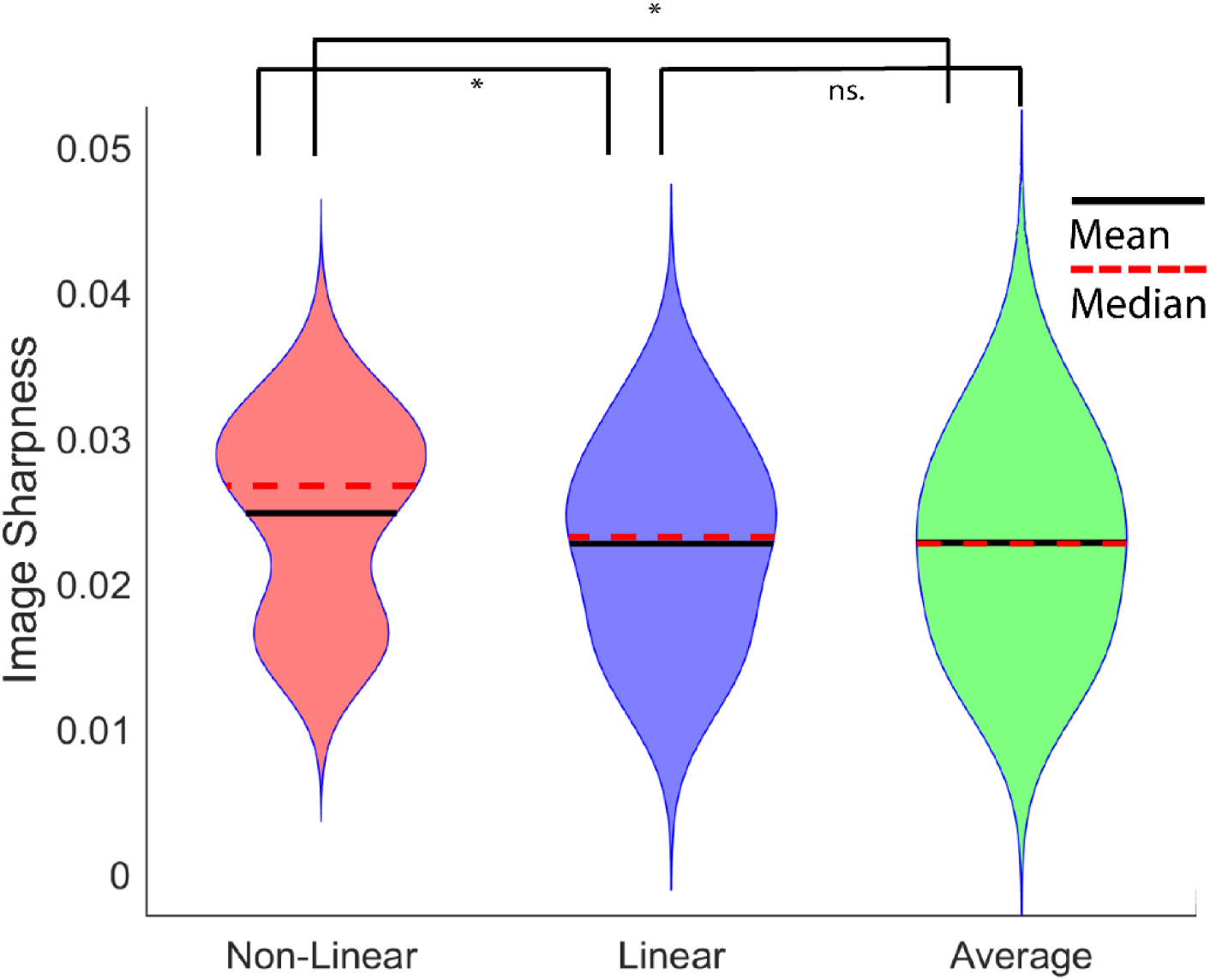
Violin plot for the three realignment methods of sharpness scores of individual subject TSEs, collapsed over the three groups. * = ps < 0.05, Wilcoxon rank sum test.

Examining group differences, it was found that MND and HC groups individually did not show significant differences in sharpness scores between any of the three methods, though the same pattern of results was found for Dice overlaps in MND and HCs individually as when we collapsed across groups (i.e., *Non-Linear* > *Linear* > *Average*). The YHP group alone showed the same (significant) pattern of sharpness and Dice overlap results as when collapsing across groups.

To assess the performance of the methods at mitigating motion artefacts, manual ratings for quality assurance were first checked for inter-rater reliability. High inter-rater reliability was found, with an overall ICC of .824, 95% confidence intervals = .778 -.861. Individual criteria ICCs were also in the high (.75+) range. We found that the MND and the older aged HCs performed worse in their motion rating scores than YHPs (Figure 5). MND and HCs had more ringing and other artefacts, worse CNR in both subcortical structures and between WM and GM boundaries, and lower sharpness. We also observed 8 YHP participants with anatomical variations including hippocampal cysts and incomplete hippocampal inversions. We observed severe neurodegeneration in MND patients.

**Figure 5.**
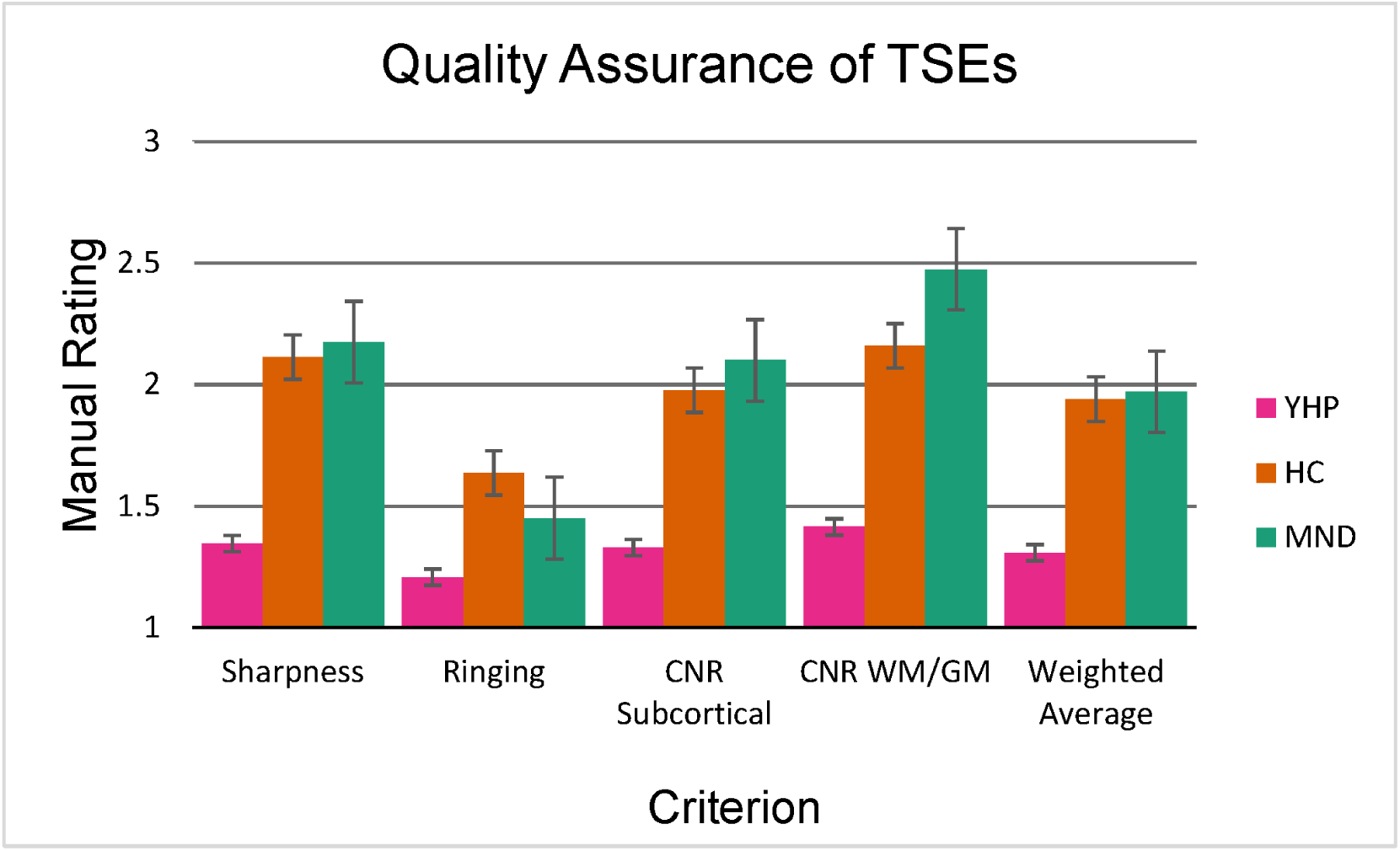
Quality assurance of TSEs for YHPs, HCs, and MND patients. Average ratings (from 1-3) with standard error of the mean are shown for the four assessed criteria and weighted average.

Interestingly, in a subset of participants (N=1 for MND, and N=8 for YHP) participants, *Linear* and/or *Average* registration techniques out-performed *Non-Linear* for Dice overlap. A case-by-case analysis of these participants was performed. In eight of these cases, there was an overall Dice failure (Dice = < 0.7) for template overlaps for all methods (i.e., *Linear* and *Average* included). We found five of the nine participants contained either anatomical variability (e.g., incomplete hippocampal inversion, cysts) that caused mislabelling in all conditions, which suggests *Linear* or *Average* performed better by chance in these participants. In the four remaining under-performing *Non-Linear* participants, we found little or no movement (e.g., in YHPs). Most of the participants in the YHP group had no adverse motion artefacts before processing. Figure 1 shows examples of labelled method templates and corresponding labelled subject TSEs.

## 4. Discussion

We investigated the influence of different realignment strategies for repeated high-resolution images of the hippocampus and found that *Non-Linear* realignment out-performs *Linear* and *Average* registrations for improving segmentation consistency for the hippocampus on a group level. This improves the robustness of brain image segmentation and mitigating the effects of motion artefacts in high-resolution anatomical MRI. We found that non-linear registration assists in registration consistency between individuals and a representative group average compared to other averaging techniques. It was hypothesised that greater registration consistency would occur between participants if the registration procedure is more effective and the between-subject registrations would consequently converge more readily due to increased boundary delineation. The results largely reflect that *Non-Linear* realignment is a suitable technique on the whole for increasing image sharpness, which leads to better segmentation consistency. We suggest that participant motion during high-resolution hippocampus acquisitions is mitigated by non-linear realignment.

Studying individual participants, *Linear* and *Average* registration techniques occasionally out-perform *Non-Linear* in participants with little or no movement, but only in YHPs. From examining these ‘failed’ cases, it is apparent that non-linear realignment may have only modest effects when there is no movement, image artefacts, or CNR issues. It was found that in most of the cases where *Non-Linear* was an under-performer in segmentation accuracy, *Average* and *Linear* techniques also failed in terms of Dice overlap. These cases are visible in Figure 3 and fall in the lower range of Dice scores. In the remaining cases where only modest benefits of non-linear realignment are observed, we propose that compounding interpolation errors during the realignment process and regularisation errors are responsible for the poor performance in these participants. Multiple non-linear registrations to an intermediate space require multiple interpolation steps in the *Non-Linear* method. It is possible that participants with minimal movement were adversely affected by (the largely unnecessary) non-linear registrations. In most participants where *Linear* or *Average* out-performed *Non-Linear*, we note the anatomical variability of five participants that influenced the segmentation of the participants. We found three incomplete hippocampal inversions, and two participants with large hippocampal cysts, which negatively impacted the segmentation consistency in all three methods. These aberrations in anatomy were not reflected in the overall templates and therefore Dice overlaps were negatively affected by aberrations in the labelling. In MND patients, we found many anatomical abnormalities. However, the MND group also showed the largest movement-related artefacts. The segmentation failures evident in this group were mitigated by the non-linear registrations.

In participants with severe motion artefacts, non-linear realignment showed its greatest utility, with much of the information lost to partial volume effects or motion artefacts being reclaimed through the technique. Older aged and diseased populations may also show decreased contrast (Wisse, Biessels, & Geerlings, 2014). Our results show consistent and improved automatic segmentation can be achieved in older aged and diseased populations using non-linear registrations, which are notoriously difficult to obtain. MND and HCs, who generally have more motion artefacts, showed the most improvement for segmentation consistency with non-linear realignment, strengthening this case. However, we found no significant differences in sharpness in the MND group between methods. This may suggest difficulties resolving SNR and contrast decreases in older age groups.

Non-linear registrations (especially ANTs SyN) incur a high computational cost. At higher resolution, these computational costs may be prohibitive. The symmetrical condition of the registration does not necessarily need to be fulfilled in order for the realignment to be successful. Therefore, other non-linear registration platforms including Greedy (Xie et al., 2018) or VolGenModel (Janke & Ullmann, 2015) should be explored in future works.

Measuring segmentation consistency is not a direct measurement of any realignment technique’s effectiveness, and the overlap resulting from registrations may not provide a true representation of segmentation *accuracy*. Here, we report on segmentation consistency, though consistency may have been more reliably derived through manual segmentations. Our measure of image sharpness offers convergent validity to the measure of image segmentation, as sharpness was found to be highest in the *Non-Linear* condition. Image sharpness largely reflects the smoothness of anatomical boundaries, with higher sharpness denoting better delineation between anatomical structures. Hippocampus segmentation relies on the distinction between anatomical landmarks such as Cornu Ammonis 1 and Dentate Gyrus (as separated by the granule cell layer). These features can clearly be seen, e.g., in Figure 2, and we are therefore confident our measure of segmentation consistency is meaningful.

We conclude that hippocampus image segmentation can benefit from non-linear registrations when participant motion is an issue. It is proposed that due to the high computational cost, non-linear realignment be used judiciously in participants with high inter-scan motion, as determined by metrics such as sharpness or qualitative assessment. In participants with no discernible motion artefacts (a rare occurrence in 12 minute acquisitions), it is suggested that linear realignment be used.

## Acknowledgements

The authors acknowledge the facilities and scientific and technical assistance of the National Imaging Facility, a National Collaborative Research Infrastructure Strategy (NCRIS) capability, at the Centre for Advanced Imaging, The University of Queensland. MB acknowledges funding from Australian Research Council Future Fellowship grant FT140100865. This research was undertaken with the assistance of resources and services from the Queensland Cyber Infrastructure Foundation (QCIF). The authors acknowledge Aiman Al Najjer for acquiring data, and the facilities at The Centre for Advanced Imaging at the University of Queensland. The data acquisition was partly funded by a study from the Cooperative Research Centre for Mental Health.

